# Machine-learning-based determination of sex-related bladder cancer biomarkers

**DOI:** 10.64898/2025.12.20.695175

**Authors:** Joseph R. Pizzi, Image Adhikari, Prakyat Prakash, Hiroshi Miyamoto, Feng Cui

## Abstract

Bladder cancer exhibits sex-specific behavior, occurring more frequently in males but progressing to advanced stages more commonly in females. The activation of sex hormone receptors may explain these differences, but the exact genetic drivers remain poorly understood. Furthermore, current bladder cancer biomarkers have inconsistent sensitivities and specificities in practice, making early diagnosis a challenge. This study approaches bladder cancer biomarker discovery through machine learning techniques on gender and disease-stratified RNA-seq data. Training sets limited to differentially expressed genes were subjected to four different feature selection methods: differential gene expression analysis adjusted p-value, recursive feature elimination with support vector machine, logistic regression, and an optimized random forest procedure. Gene panels were compared and aggregated across selection strategies and cross-validation folds to identify robust biomarkers for sex-specific bladder cancer development and progression. When applied to unseen datasets and limited to 50 genes or less, male and female-specific panels achieved areas under the receiver operating characteristic curve of 0.932 and 0.914, respectively, in distinguishing bladder cancer samples from non-tumor controls. Genes such as PRAC1 and PCDH11Y were identified as high-impact predictors related to sex hormones or chromosomes for male tumor development. In the female-specific panel, genes related to aberrant androgen signaling across tumor types like AR, PLXNA1, USP54, and PMEPA1 were influential. These results offer potential targets for further in vivo/vitro experimentation and provide a framework for constructing generalizable, high-performance gene panels for bladder cancer diagnosis and prognosis.

## Introduction

Bladder cancer (BCa) is the 6th most common cancer in the United States, representing 4.2% of all cancer cases (1). The 5-year relative survival rate is estimated at 79.0%, but individual prognosis depends heavily on the stage and aggressiveness of the tumor. However, identifying BCa early in development can be difficult, with most symptomatic individuals presenting with gross hematuria, a malady common in other diseases like urinary tract and kidney infections (2). The current diagnostic standard is cystoscopy, yet this method relies on human judgment and can miss carcinoma in situ without further imaging (3). Cystoscopy is often preceded by urine cytology to identify abnormal cells characteristic of high-grade tumors but this technique can prove ineffective for lower-grade cases. To mitigate risk while prioritizing prompt detection, the FDA has explored and approved less-invasive molecular biomarkers for use in the clinic. Unfortunately, their performance is unreliable, with nuclear matrix protein 22 and bladder tumor antigen tests exhibiting highly variable sensitivities and specificities (4). While they offer an improvement upon cytology in terms of sensitivity, current BCa biomarkers still suffer from high false-positive rates.

Gender-specific differences in frequency and pathogenesis add another layer of complexity to the task of better diagnosing and treating BCa. Across all races, the incidence rate of BCa in males is approximately 3.3 times what is observed in females (5). Only part of this phenomenon can be attributed to a higher percentage of men being smokers, with a 2020 study finding that men had a prevalence of 32.6% while women stood at 6.5% (6). Despite the stark difference in frequency, females tend to have more aggressive forms of BCa upon diagnosis. A recent French study found that only 8.9% of women with muscle-invasive BCa had a history of the non-invasive variant, while men did at a higher rate of 26% (7). Systematic biases in the healthcare system can partially explain this. Women presenting with hematuria are typically less likely to be referred to a urologist, with a 29% referral rate compared to 45% for men in NCI’s Southern Community Cohort Study (8). However, females still tend to have lower survival rates compared to men, even after correcting for confounding variables like tumor stage and demographics (9).

These findings suggest that there are molecular mechanisms driving gender-specific differences in BCa outcomes in addition to population characteristics. Differences in androgen receptor (AR) signaling have been explored as a critical factor in BCa sex dimorphism (10). In canonical signaling, the binding of androgens like testosterone and dihydrotestosterone to AR forms a dimer complex that reaches the nucleus and binds androgen response elements to shift transcriptional patterns (11). In bladder tissue, AR signaling has been linked to urine storage, urinary tract function, nerve functions, and urothelium musculature/thickness (10). Preliminary research suggests that aberrant AR activity may influence BCa development/progression through potential downstream targets like ADGRL3, ELK1, and GABBR2 (12–14). Individuals with male reproductive organs tend to have higher levels of androgens in their blood circulation than females, which could contribute to the increased prevalence of the disease in men (15). On the other hand, estrogen receptor (ER) activity in females may contribute to the observed increased tumor aggressiveness compared to males. In traditional ER signaling, similarly to the mechanism of AR activity, estrogen-bound ERα or ERβs act as transcription factors (16). Reviews of current BCa research have revealed the importance of ERβ, with ERβ positivity ranging from 27-100% in urothelial tumors across various publications compared to 0-38% exhibiting ERα positivity (17). Potential downstream effectors of ERβ in BCa cells have been identified. For example, ERβ activation was demonstrated to reduce GULP1 expression to elicit cisplatin resistance in BCa cell lines (18). Outside of steroid hormone signaling, sex chromosome differences may contribute to BCa sex dimorphism as well. For example, the KDM6A gene is a tumor suppressor located on the X chromosome that can escape inactivation, leading to enhanced protection in females due to having an extra copy (19). Specifically, KDM6A-lacking organoids implanted in mice had increased expression of basal genes and decreased expression of luminal genes (20). Overall, the genetic and cellular mechanisms underlying higher BCa prevalence in men but more aggressive progression in women are still unclear and require further study.

The lack of robust, sensitive biomarkers in tandem with established gender differences in disease development calls for the discovery of new BCa biomarkers. This study approached this problem using machine learning, where expression data is split into training and testing sets for a classification task. Here, three classification tasks were handled with machine learning: male/female tumor versus non-tumor and male versus female tumors based on publicly available tissue-level high-throughput sequencing data. First, differential gene expression analysis (DGEA) was applied as a filter-based feature selection method to reduce the sparsity of the gene expression data and highlight differentially expressed genes (DEGs) related to sex and tumorigenesis. Then, four different feature selection approaches - DGEA adjusted p-value, an optimized random forest (RF) procedure, recursive feature elimination with support vector machine (SVM-RFE), and logistic regression - were applied to task-specific DEGs to isolate high-impact biomarkers. Afterward, the rankings for each approach were aggregated across fold schemes into one synthesized gene panel for each classification task using robust rank aggregation (RRA). The effectiveness of aggregated and single-method panels was evaluated on external testing sets using multiple machine learning models based on a combined metric that takes class imbalance into account. Pathway enrichment and protein-protein interaction (PPI) network analysis were also carried out to reveal the prominent molecular mechanisms within each aggregated panel. In the end, robust, biologically relevant, and sex-related biomarkers for BCa development and progression that generalize well to unseen data were identified.

## Results

### Dataset Merging

In classifying male tumors versus female tumors, the TCGA-BLCA dataset had a reasonable number of samples for each group: 267 male tumors and 102 female tumors after removing stage I, metastasized, and low-grade samples from consideration (Table 1). However, under 5% of the entire TCGA dataset consisted of non-tumor controls, making the sex-stratified tumor versus non-tumor task nearly impossible to execute. In response, a merged cohort was created to combat this extreme class imbalance. TCGA-BLCA, GTEx, and GSE133624 were independently processed and z-scored before being concatenated together on common genes in the GRCh38.p13 reference genome. To visualize the effects of z-score merging, Uniform Manifold Approximation and Projection (UMAP) dimensionality reduction was performed on the normalized transcripts per million (TPM) data (Figure 1). When the three datasets were merged without independent z-scoring, samples clustered strictly based on their dataset. When z-scoring before concatenation, this clustering pattern was disrupted, and a more heterogeneous mixture of samples across datasets was observed.

**Figure 1:**
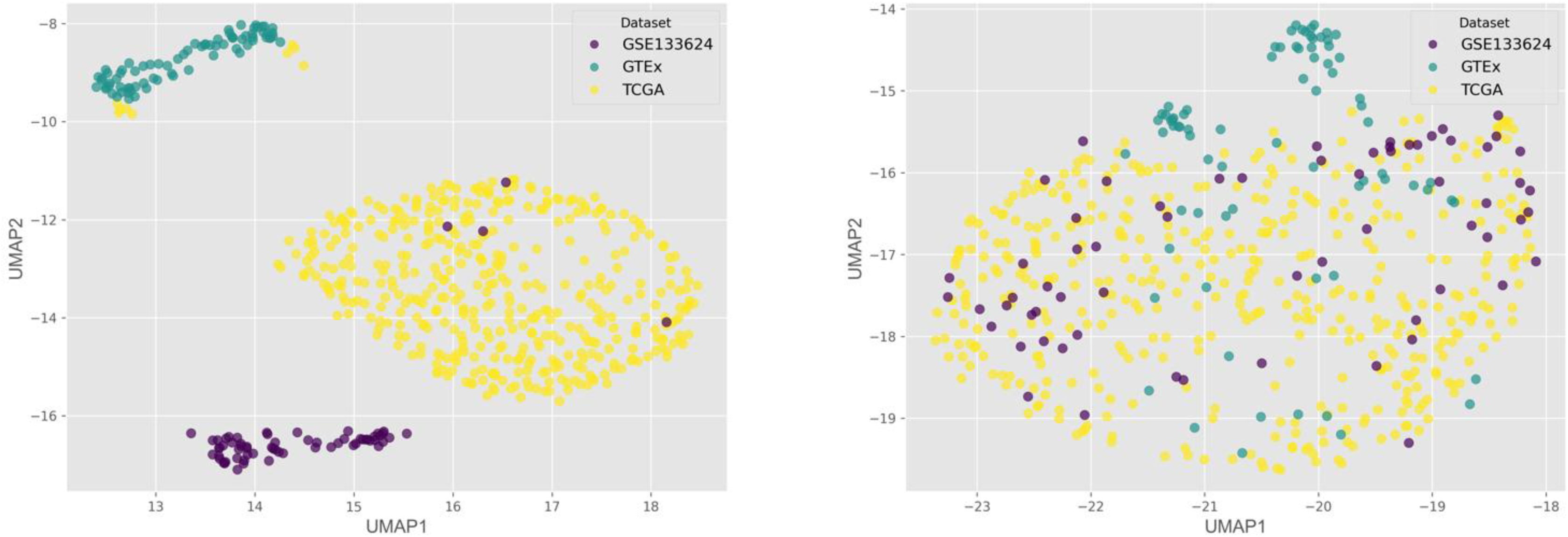
UMAP projections of merged dataset normalized TPM values before (left) and after (right) implementing independent z-scoring prior to merging.

**Table 1:** Class distribution of each dataset. TCGA, GTEx, and GSE133624 were independently z-scored, then merged to constitute the training set for the male/female non-tumor versus tumor analyses. When used alone, the TCGA data was corrected to remove metastasized and lower staged samples.

| Dataset | Use | Total | Male Tumor | Female Tumor | Male Non-Tumor | Female Non-Tumor |
| --- | --- | --- | --- | --- | --- | --- |
| TCGA | Training | 431 | 304 | 108 | 10 | 9 |
| GTEX | Training | 77 | 0 | 0 | 48 | 29 |
| GSE133624 | Training | 65 | 32 | 4 | 25 | 4 |
| GSE236932 | External | 58 | 27 | 8 | 15 | 8 |
| GSE188715 | External | 70 | 46 | 11 | 11 | 2 |

### Differential Gene Expression Analysis and Feature Selection

Within each fold scheme, DGEA was carried out for male/female non-tumor versus tumor samples and male tumors versus female tumors. For all analyses, DEGs were defined as those with an adjusted p-value less than 0.05 and a log2 fold change greater than |1.0|. After comparing and excluding overlapping genes across cohorts, the result was three groups of DEGs: male-specific BCa development, female-specific BCa development, and sex-related BCa progression. This was also performed outside of the cross-validation scheme on the entire training dataset for each task and visualized with volcano plots (Figure 2). For the male-specific tumor development analysis, 2800 DEGs were identified with 1414 upregulated and 1386 downregulated. In the female-specific development task, only 821 DEGs were significant with 525 upregulated and 296 downregulated. Finally, in comparing expressions between male and female samples, 328 DEGs were exclusive to diseased tissue with 223 upregulated and 105 downregulated.

**Figure 2:**
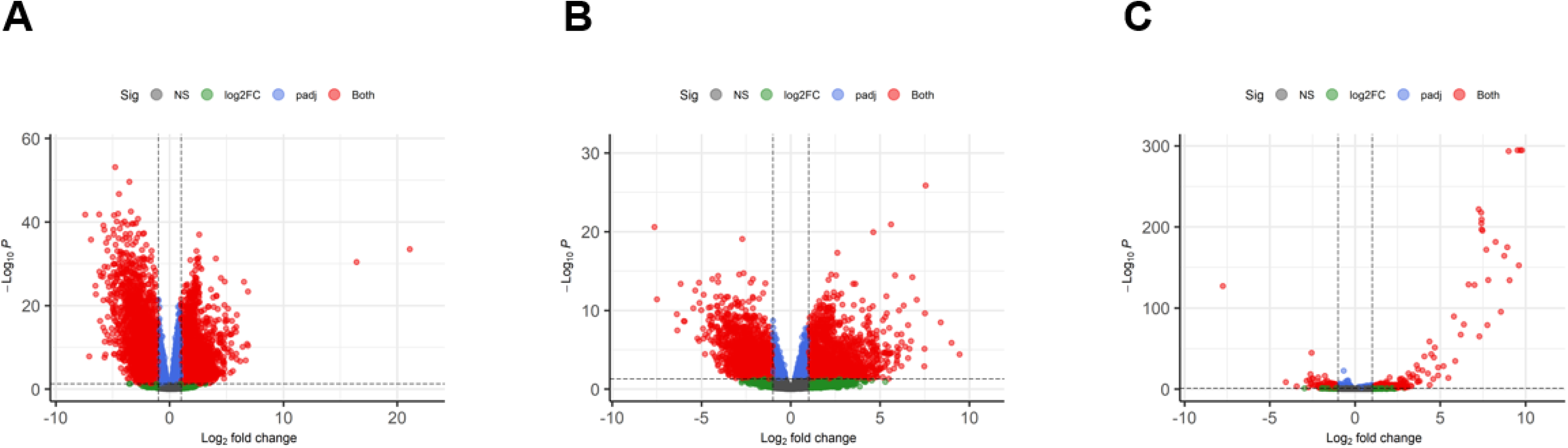
Volcano plots of DGEA for male tumor versus non-tumor (A), female tumor versus non-tumor (B), and male versus female tumors (C).

Each DEG pool was then subjected to feature selection using four different methods: optimized RF, SVM-RFE, logistic regression, and filtering via the DGEA adjusted p-value. Each strategy isolated the top 300 DEGs based on composite metric performance, which combined F1 score, the area under the receiver operating characteristic curve (AUROC), and balanced accuracy to account for class imbalance. Each task produced four top 300 gene panels in five different fold schemes to create 20 rankings in total. Through RRA, these were consolidated into an aggregated top 300 ranking, where genes placing highly across all 20 input panels were associated with a better score (Table S1-S3). The top 20 genes for each task in terms of RRA score were visualized as a bar graph by transforming their scores with a negative log transformation (Figure 3A). All cohorts had a clear top gene: NDNF, ARSF, and LINC03020 for male, female, and tumor, respectively, with scores leveling off towards the end of the top 20. The top 300 rankings for each selection method were also aggregated across fold schemes to create cross-fold consensus rankings, which were then compared to each other with a Venn diagram (Figure 3B). Regardless of the classification task, optimized RF and SVM-RFE tended to produce quite distinct panels, while logistic regression and DGEA adjusted p-value strategies had more genes in common with other methods. The male non-tumor versus tumor task had the least amount of agreement between the four selection methods, with only four genes being shared among all the consensus panels. On the other hand, different selection methods had much more of an overlap in the male tumor vs. female tumor task, with 135 genes shared between the aggregated panels for each.

**Figure 3:**
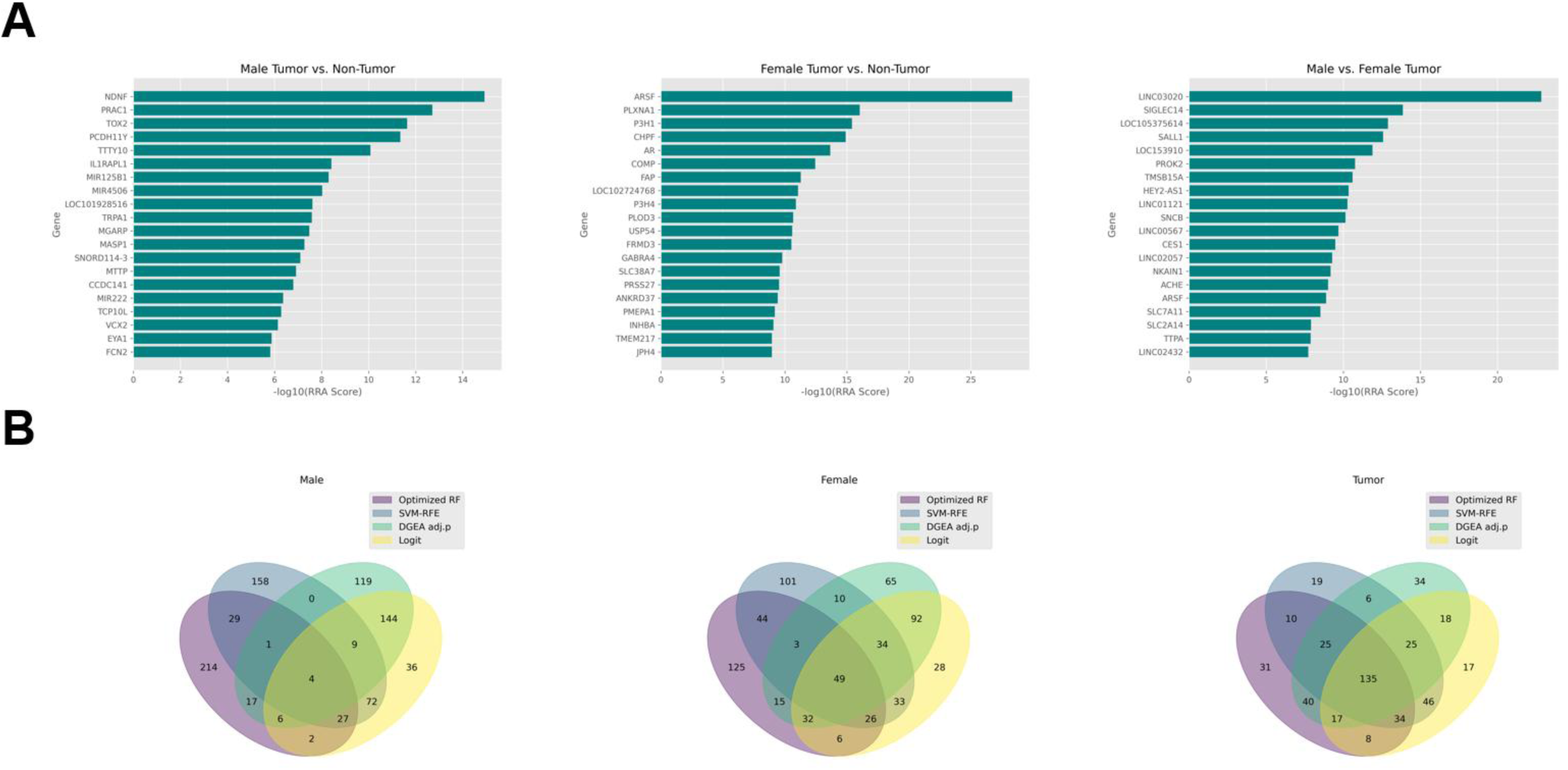
Aggregation of gene panels across fold schemes and selection methods. (A) Bar graphs depicting the top 20 genes in terms of RRA score in the cross-fold cross-method aggregated panel for each classification task. A negative log10 transformation was applied to each score. (B) Venn diagrams displaying the overlap in genes between the cross-fold consensus panels for each selection method in each classification task.

### Internal Validation

In each fold scheme, the testing fold acted as an internal validation set to determine how well the selected genes applied to each classification task on data similar to the training data. Four machine learning models, RF, SVM, K-nearest neighbors (KNN), and XGBoost, were hyperparameter optimized, fit on the four training folds, then tested on the final fold. Balanced accuracy, F1 score, AUROC, and the composite metric were averaged across all 5 folds for a given gene panel in all three classification tasks. The aggregate panels were not considered, as those were assembled outside of the cross-validation scheme. In the male tumor versus non-tumor classification task, the best performing panel was the optimized RF top 50 using XGBoost as the evaluation model with a composite metric of 0.988 (Table S4). The following four best panels were also all derived from the optimized RF top 300, but some used RF as the evaluation model. In the female tumor versus non-tumor analysis, the top two panels were the SVM-RFE top 300 and 200 employed with XGBoost, demonstrating average composite scores of 0.965 and 0.963, respectively (Table S5). Comparatively, the male tumor versus female tumor classification was much more difficult, with the logistic regression top 300 using RF evaluation having the best average composite metric of 0.734, but a more respectable average F1 score of 0.843 (Table S6).

### External Validation

The same four models used for internal validation were re-trained on their respective full, unfolded training dataset then applied to two external datasets, GSE236932 and GSE188715. The cross-fold cross-method aggregated panel and the cross-fold panels for each selection method were considered in each classification task. First, ROC curves were constructed to compare between the aggregated and cross-fold individual selection method panels in each task/dataset combination (Figure 4). Due to their consistent high performance in internal validation, the SVM-RFE and Optimized RF panels with the highest AUROC were chosen to represent the individual selection methods. In the male tumor versus non-tumor classification, the AUROC of the best aggregated panel surpassed those of the single selection methods in the first external dataset and was comparable in the second. For female samples, the top aggregated panel was within 0.1 of the optimized RF selection AUROC in GSE236932, and all panels were tied at a perfect 1.0 due to sample size limitations in GSE188715. Finally, in differentiating male tumors from female tumors, the aggregated panel had the highest AUROC in both external datasets. In GSE236932, SVM tended to be the best-suited evaluation model across all gene panels, but this shifted to XGBoost and RF in GSE188715. Across all datasets and panels, the highest AUROCs were obtained by panels with the top 100-300 genes. To examine the aggregated panels further, the performance metrics of each panel were averaged across both external datasets (Table S7-S9). Then, the best aggregated panels with 50 or less genes were considered (Table 3). The aggregated male top 50 was able to achieve average AUROC and F1 scores of 0.923 and 0.917 (0.915 composite), respectively with RF. In the female classification task, the top 10 genes in the aggregated panel produced slightly poorer, but still respectable scores of 0.914 and 0.878 for AUROC and F1 (0.863 composite), also using RF as the evaluation model. A notable decline in performance was observed in the male tumor versus female tumor task, with the aggregated top 25 genes yielding a composite score of 0.784. In all tasks besides male tumor versus non-tumor, the best aggregated panel with less than 50 genes had a balanced accuracy less than 80%.

**Figure 4:**
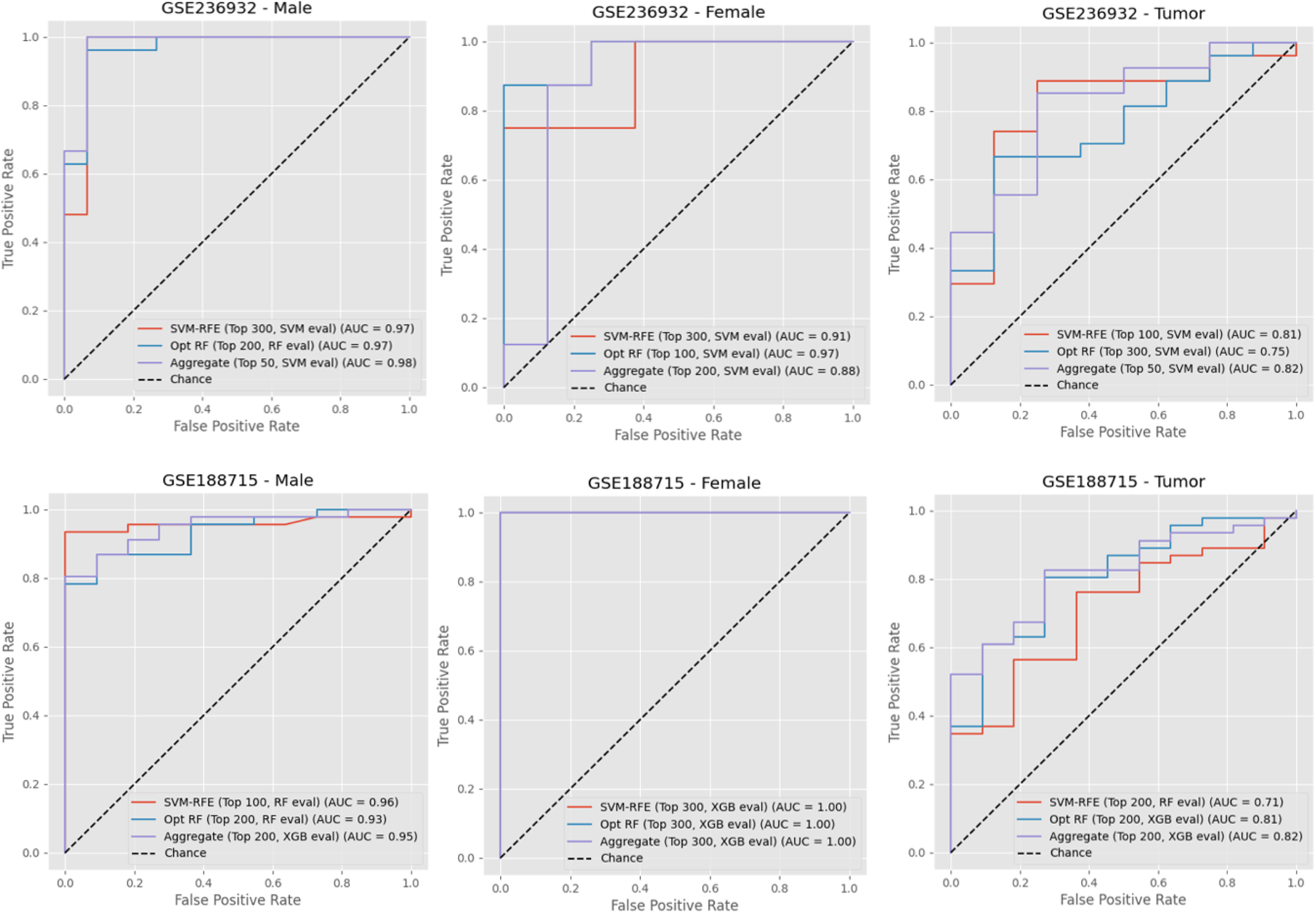
ROC curves of external validation performance. The panels with the highest AUROCs for the optimized RF, SVM-RFE, and aggregated rankings were chosen for visualization in each combination of classification task and external dataset. SVM-RFE and optimized RF were selected based on internal validation performance.

### Pathway Enrichment, PPI Network Analysis, and Literature Search

To determine the pathways and biological processes relevant to the machine learning-selected genes, enrichment analysis with GO (Figure 5A) and KEGG (Figure 5B) databases was performed. Only terms with a q-value of less than 0.05 were considered. When the aggregated top 300 panels were analyzed, only the male tumor vs. female tumor task yielded significant enrichments. The top ontologies were retinol metabolism and hormone metabolic process for KEGG pathways and GO biological process, respectively. Relevant KEGG terms included the metabolism of drugs and xenobiotics by cytochrome P450, chemical carcinogenesis, and steroid hormone biosynthesis. In terms of GO biological process enrichments, xenobiotic response and metabolism, estrogen metabolic process, and cellular glucuronidation were especially pertinent. In order to get enrichments for the sex-specific panels, the search was expanded to include the top 500 genes of the aggregated panel. For male tumor versus non-tumor, the top terms were cGMP-PKG signaling and cell-substrate adhesion for KEGG and GO, respectively. Other tumor-related ontologies included cytoskeleton in muscle cells, focal adhesion, hormone catabolic process, and junction assembly. For female tumors versus non-tumor, the terms with the lowest q-value, oddly, were cornified envelope formation and skin development for KEGG and GO. However, some terms more relevant to the tumor context also appeared, like extracellular matrix organization, collagen fibril organization, and protease inhibition.

**Figure 5:**
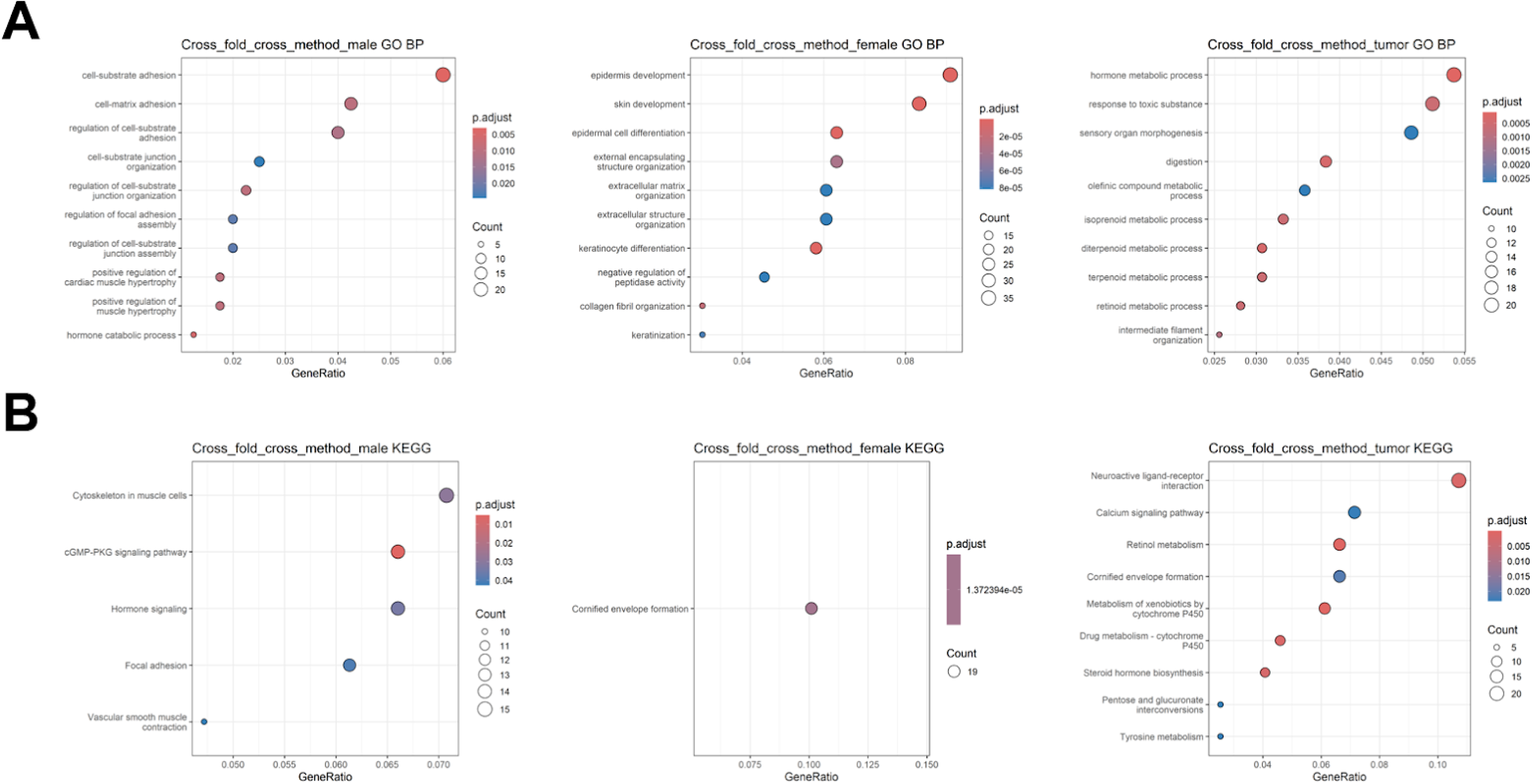
Dot plots for pathway enrichment analysis. (A) Top 10 enriched terms for the GO biological process database in terms of q-value and gene ratio (# of genes associated with pathway : # of genes) for each classification task (B) Top 10 enriched terms for the KEGG pathway database, but the male and female tumor versus non-tumor top 500 genes yielded less than 10 significant enrichments.

A PPI network was also constructed for each of the aggregated gene panels corresponding to the three classification tasks. In the male panel, ESPL1, NUSAP1, KIF22, ZWILCH, IGF1, PTTG1, PDGFRA, FGF9, MCM7, and DKC1 were the top 10 hub genes based on maximum clique centrality. Two significant gene clusters contained hub genes (Figure 6A). The first contained PTTG1, KIF22, ZWILCH, NUSAP1, and ESPL and was related to sister chromatid separation and mitotic anaphase based on Reactome pathway enrichment. The second, comprised of PDGFRA, BDNF, FGF19, IGF1, and FGF9, was associated with aberrant PI3K-AKT signaling cancer. For the female cohort, COL5A2, PLOD1, COL12A1, P3H1, COL2A1, BGN, SERPINH1, COLGALT1, PLOD3, and P3H4 were hub genes. One significant gene cluster contained all female hub genes, and was significantly enriched for collagen formation and extracellular matrix organization (Figure 6B). Finally, the hub genes for the male tumor versus female tumor network were NTRK2, CALB1, UGT2B15, TH, CYP1A2, GRIN2B, ADH7, WNT3A, WNT10B, and WNT16. Three significant gene modules contained hub genes (Figure 6C). The first cluster was made up of CALB1, PPP1R1B, TH, GRIN2B, and NTRK2, and was connected to NTRK signaling. The second, containing WNT10B, SOST, CCN4, WNT3A, and WNT16, was primarily related to WNT signaling and ligand biogenesis. The third cluster was the largest and consisted of AKR1B10, SULT1E1, CYP1A2, UGT2B15, ALDH3A1, CES1, CYP26B1, and ADH7. The enrichments for this final cluster were vague, with biological oxidations and phase I/II of metabolism.

**Figure 6:**
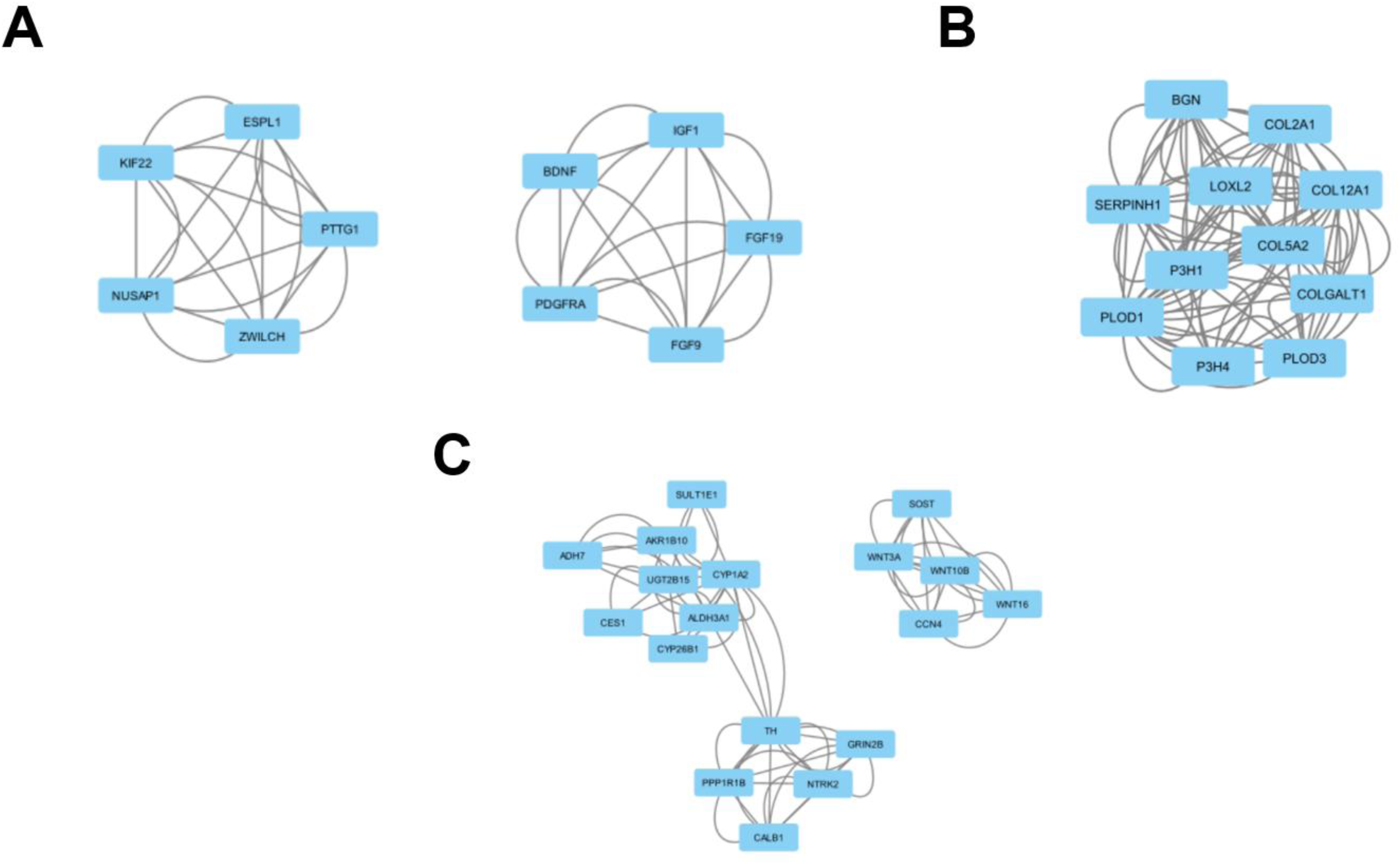
Significant gene modules containing hub genes in PPI network analysis for male (A), female (B), and tumor (C) cohorts. Hub genes were defined as genes in the top 10 maximum clique centrality ranking while significant gene modules were defined as having a score of 3.00 or greater.

Due to the difficulties in obtaining relevant enrichments, a detailed literature search was also performed for the top 20 genes in each aggregated panel (Table 2). Genes were categorized as BCa-related, tumor-related, or other. BCa-related genes had to be connected to BCa either through bioinformatics analyses or in vitro/vivo experiments, while tumor-related genes could be for any tissue type. Both the sex-specific development panels contained more BCa-relevant genes compared to the sex-related progression selection. The latter contained many more poorly characterized lncRNAs. In the male-specific panel, PRAC1 was of particular interest due to its high ranking and connection to prostate cancer. Additionally, sex chromosome-linked genes like PCDH11Y, TTTY10, and IL1RAPL1 appeared as strong predictors of male tumor development. On the female side, many genes related to androgen signaling, like AR, USP54, PMEPA1, and PLXNA1 were in the top 20 of the aggregated panel.

**Table 2:** Literature search of genes in the top 20 of the aggregated panel based on RRA score for each classification task. BCa-related genes appeared in either published bioinformatics analyses or in vitro/vivo experiments of BCa. Many of these were also relevant to other tumor types.

| Task | BCa-related | Tumor-related | Other |
| --- | --- | --- | --- |
| Male Tumor vs. Non-Tumor | NDNF, PRAC1, TOX2, TRPA1, SNORD114-3, MTTP, CCDC141, MIR222 | PCDH11Y, TTTY10, IL1RAPL1, MIR125B1, MASP1, TCP10L, EYA1, FCN2 | MIR4506, LOC101928516, MGARP, VCX2 |
| Female Tumor vs. Non-Tumor | P3H1, CHPF, AR, FAP, P3H4, PLOD3, USP54, PRSS27, ANKRD37, PMEPA1, INHBA | ARSF, PLXNA1, COMP, FRMD3, GABRA4, SLC38A7, TMEM217, JPH4 | LOC102724768 |
| Male Tumor vs. Female Tumor | SNCB, SLC7A11, SLC2A14 | SIGLEC14, SALL1, LOC153910, PROK2, TMSB15A, LINC01121, LINC00567, CES1, LINC02057, NKAIN1, ACHE, ARSF, TTPA, LINC02432 | LINC03020, LOC105375614, HEY2-AS1 |

**Table 3:**
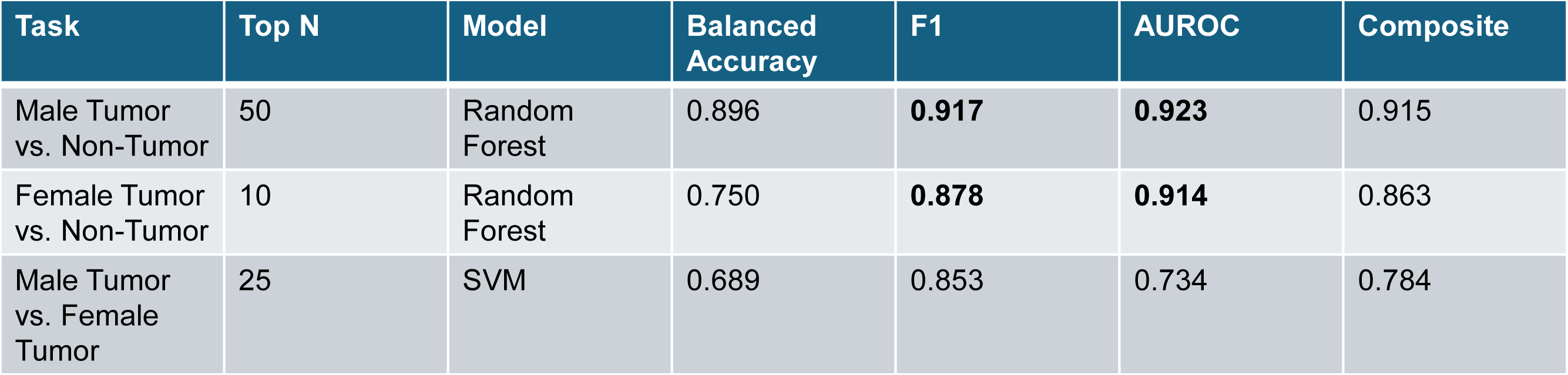
Best performing aggregate panels with n ≤ 50 for each classification task. For each aggregate panel, the number of genes, evaluation model, and average performance metrics across both external datasets are provided. The composite score was weighted to favor metrics better suited for imbalanced datasets: 0.5*F1 + 0.3*AUROC + 0.2*balanced accuracy.

| Task | Top N | Model | Balanced Accuracy | F1 | AUROC | Composite |
| --- | --- | --- | --- | --- | --- | --- |
| Male Tumor vs. Non-Tumor | 50 | Random Forest | 0.896 | **0.917** | **0.923** | 0.915 |
| Female Tumor vs. Non-Tumor | 10 | Random Forest | 0.750 | **0.878** | **0.914** | 0.863 |
| Male Tumor vs. Female Tumor | 25 | SVM | 0.689 | 0.853 | 0.734 | 0.784 |

## Discussion

This study adopts a novel approach in identifying sex-related BCa genes and highlights new candidate biomarkers for further experimentation. A recent study by Wang and colleagues highlighted sex-specific genes through DGEA, PPI network construction, pathway enrichment analysis, and immune cell infiltration correlation (21). Here, DGEA was the starting point for a rigorous cross-validated machine learning workflow using four different feature selection techniques. Distinct sets of genes were generated by each selection method, therefore an ensemble-like approach with RRA was employed to generate robust, high-performing biomarker panels. In external validation, male and female-specific gene panels aggregated across fold schemes and feature selection techniques limited to 50 or fewer genes obtained average AUROC scores of 0.932 and 0.914, respectively. These models achieving AUROC values greater than 0.90 indicate a respectable diagnostic performance in the context of clinical predictions (22). Pathway enrichment and PPI analysis confirmed the molecular pathways associated with carcinogenesis in each sex, with males displaying a cell interaction, mitotic regulation, and PI3K-AKT-focused signature while the female panel consisted of ECM organization and collagen synthesis-connected genes. A literature search of the genes in each sex-specific BCa development panel revealed that 4-5 top predictors were tumor-related and either sex-linked or associated with sex hormone signaling in both. In contrast, the male versus female tumor classification task displayed much poorer performance in both internal and external validation. Even in the best-performing aggregated panel, an average composite score of 0.784 with an average AUROC of 0.734 was obtained in external validation. However, a literature search showed that many of the top genes were still relevant to tumors and further analysis found that genes involved in xenobiotic metabolism were prominent in the aggregated panel.

For the male-specific tumor development genes, PRAC1, TTTY10, PCDH11Y, and IL1RAPL1 were connected to sex-related chromosomes or molecular mechanisms. Alternative names for PRAC1 include prostate cancer susceptibility candidate protein 1 and prostate, rectum and colon expressed gene protein. Previous studies have found that PRAC1 loss in prostate cancer led to cell growth even in AR signaling inhibitor-treated samples and may act as a co-regulator of AR activity (23). In the context of BCa, PRAC1 methylation was shown to be significantly higher in BCa tissue and associated with higher tumor grade, as well as demonstrating significant downregulation in bioinformatics analyses (24–26). Interestingly, PCDH11Y, a Y-linked protein-coding gene, was identified as having increased mRNA expression in androgen-resistant prostate cancer cell lines (27). TTTY10, a Y-linked lncRNA, has been implicated in numerous tumor types, including colorectal cancer, papillary thyroid carcinoma, osteosarcoma, and esophageal squamous cell carcinoma (28–31). Finally, IL1RAPL1 is an X-linked gene thought to be involved in the pro-inflammatory response through NF-kB and AP-1 pathways that is overexpressed in tumors like pancreatic ductal adenocarcinoma, Ewing sarcoma, and triple-negative breast cancer (32). Outside of genes connected to sex-specific chromosomes or pathologies, MIR222 was also in the top 20 of the aggregated panel. Functional analyses have identified that METTL3 promotes BCa proliferation by maturing pri-MIR221/222, which then go on to target PTEN (33). Other genes like NDNF, TOX2, TRPA1, SNORD114-3, and MTTP have been noted as significantly downregulated, upregulated, and/or associated with pathological features in BCa compared to controls (25,34–37). Moreover, when performing pathway analysis on the top 500 genes in the aggregated panel, numerous tumor-relevant terms were significantly enriched. Wnt/β-catenin transcription has been linked to cGMP-PKG signaling (the top KEGG enrichment), leading to cancer cell expansion and immune evasion (38). Specifically, in epithelial ovarian cancer, PKG activity was shown to alter EGF-elicited cell proliferation and migration (39). Moreover, focal adhesions are known to be connected to the tumor microenvironment, acting as the physical support connecting tumor cells to the surrounding ECM and allowing for mechanical signaling (40). Another significantly enriched term regarding cell interactions was junction assembly. The epithelial-to-mesenchymal transition, a prominent cancer hallmark, is known to disrupt the adhesive and signaling functionalities of many types of cell junctions (41). Moreover, PPI network analysis revealed hub gene-containing clusters associated with mitotic anaphase and PI3K-AKT signaling. Research has found that the genes responsible for sister chromatid adhesion are often underregulated in cancer, leading to chromosomal instability and poorer patient outcomes (42). The top ranked hub gene in the mitosis-related module and entire male-specific PPI network, ESPL1, was found to be involved in enhanced cisplatin resistance via the JAK2/STAT3 pathway in BCa tissues and cells (43). Hyperactive PI3K-AKT signaling is attributed to numerous cancer types, but in BCa it has been associated with the epithelial-to-mesenchymal transition specifically (44). While multiple key predictors in the male-specific aggregated panel were associated with androgen signaling and sex-linked tumor development, as a whole, the panel had strong themes of cell interactions, mitotic dysfunction, and aberrant PI3K-AKT signaling.

Looking at female-specific BCa development, AR, PLXNA1, USP54, PMEPA1, and ARSF were of particular interest. AR is often downregulated in BCa, and lower rates of AR positivity have been reported in high-grade cases compared to lower grades (45). Furthermore, AR is known to be an independent prognosticator in females, but not in males (46). PLXNA1 has not been connected to BCa in the literature, but has been shown to alter AR inhibitor effectiveness in prostate cancer, with elevated expression enabling tumor proliferation even under enzalutamide treatment conditions (47). Additionally, USP54 has displayed higher expression in prostate cancer and was correlated with AR signaling levels as well as heightened proliferation compared to silenced models (48). PMEPA1 (prostate transmembrane protein, androgen induced 1) silencing has been linked to significantly decreased BCa proliferation, migration, and invasion in vitro (49). These genes may be related to aberrant androgen signaling leading to tumorigenesis in female BCa. Finally, ARSF is an X-linked gene that has not been connected to BCa, but was associated with overall survival in glioblastoma in one study (50). In terms of significantly enriched pathways for the entire female-specific aggregated panel, several were related to extracellular matrix and collagen fibril organization. Compositional changes in the ECM are characteristic of tumor development, with tumors exhibiting greater stiffness due to upregulated collagen deposition by cancer-associated fibroblasts (CAFs) (51). Tumor-associated macrophages and CAFs in tandem influence collagen remodeling to produce a shift in fibril orientation, resulting in heightened cancer cell migration (52). This was supported by PPI network analysis as well, where the significant gene cluster containing all hub genes was found to be related to collagen formation and biosynthesis. The top hub gene among the female-specific genes was COL5A2. In bioinformatics analyses, BCa patients with lower COL5A2 expression had better outcomes for tumor grade, migration, and overall survival (53). However, unlike the male pathway enrichment, several seemingly unrelated ontologies, like cornified envelope formation and epidermis development, were highly enriched.

While the sex-related tumor progression classification demonstrated worse predictive performance, many of the top predictors were still related to BCa or other tumor types. ARSF and TMSB15A, both X-linked genes, were in the top 20 of the aggregated gene panel. As previously mentioned, ARSF is linked to tumor progression in other tumor types but has yet to be connected to BCa. On the other hand, TMSB15A has been linked to breast cancer treatment response (54). Several other non-sex-linked predictors were related to tumor-induced shifts in expression or cell behavior in BCa. Two solute carrier family genes, SLC7A11 and SLC2A14, have demonstrated aberrant expression in BCa (55). Specifically, SLC7A11 upregulation inhibited ferroptosis and promoted BCa progression in an in vitro/vivo study (56). Significant enrichments for the aggregated male tumor versus female tumor panel were centered around steroid hormones and xenobiotic metabolism/response. Recent research has shown that certain variants of xenobiotic metabolism genes are associated with higher-grade tumors or recurrence in BCa (57). Furthermore, hormone metabolic process was among the top 10 most highly enriched GO biological process terms, but estrogen metabolic process specifically also had a q-value less than 0.05. In numerous in vitro studies, estrogens have proven to accelerate cell proliferation, potentially through phospho-ERK expression, as well as downregulate tumor suppressor genes in BCa cells (17). Additionally, network analysis revealed a significant gene module associated with WNT signaling. WNT signaling has been implicated in multiple tumor types including BCa due to its activation of PI3K-AKT signaling, which then induces therapeutic resistance and a proliferative phenotype (58).

These results diverge from what Wang and company recently discovered through their analyses. Wang’s study identified the AR signaling pathway as enriched in male-specific hub genes through PPI network analysis, but here, AR expression was a strong predictor of female-specific BCa development (21). This is in line with the work of Sikic and colleagues identifying the gene as an independent prognosticator in females (46). Instead, Wang and coworkers found the Wnt signaling pathway to be highly enriched in female-specific hub genes. Out of the 300 genes in the machine learning-generated male-specific BCa development panel, only 6 genes overlapped with the male-specific hub genes identified in the Wang paper: DDX11, PALB2, ESPL1, CDH17, PDGFRA, and TGFBR2. No overlap was observed in the female-specific genes between this study and theirs. However, this alternative approach treating biomarker discovery as a feature selection problem still produced high-performance, biologically relevant findings. The majority of the top predictors in both male and female-specific aggregated panels were related to BCa or other tumor types. Genes with a heavy influence on male tumor classification across folds like PRAC1, PCDH11Y, TTTY10, and IL1RAPL1 were linked to androgen signaling in other tumor types or associated with sex-linked chromosomes. In the female analysis, AR, PLXNA1, USP54, PMEPA1, and ARSF were still implicated in androgen activity, but suggest that different downstream pathways are affected compared to males.

Pathway enrichment of both aggregated panels found that male-specific genes were related to cell adhesion and junctions while female-specific genes were more relevant to ECM and collagen organization. Finally, taking the respectable performance of the aggregated male and female-specific BCa development gene panels on external datasets into account, the aforementioned high-impact predictor genes identified in this study could be promising candidates for further experimentation in vitro/vivo.

This study has limitations to take into account as well. Due to the established difference in BCa occurrence between sexes, the datasets used for this analysis had limited female samples and therefore exhibited class imbalance. This was addressed through the use of a hybrid scoring system comprised of metrics sensitive to imbalanced datasets for hyperparameter optimization and feature selection. Moreover, there was a notable drop in performance when going from the sex-specific BCa development panels to the sex-related BCa progression panel for both internal and external validation. Therefore, more weight was assigned to the sex-specific biomarkers in terms of their potential for further experimentation. More complex models integrating mutation or clinical data in addition to gene expression may be necessary to understand biomarkers of sex-related BCa progression. Finally, while the approach has been adopted in several machine learning-based studies of gene expression in cancer, independent z-scoring may not fully address the batch effects between different datasets and sequencing technologies. Still, the models trained on the merged dataset still generalized well to external datasets after they had also been z-scored independently.

## Conclusions

The present study has generated high-performance models for male and female-specific BCa development based on RNA-seq gene expression data, achieving AUROCs of 0.923 and 0.914 on external cohorts, respectively. A literature search of the top predictors in each model revealed connections to BCa and other tumors types. Specifically, for males, PRAC1, PCDH11Y, TTTY10, and IL1RAPL1 were identified as targets for further experimentation while AR, PLXNA1, USP54, PMEPA1, and ARSF were highlighted for females. These findings provide candidate genes potentially driving the sex-specific mechanisms of BCa development while also detailing a rigorous, generalizable machine learning workflow for biomarker discovery. Analyses of male tumors versus female tumors yielded a less-impressive AUROC of 0.734. Future work will focus on adapting the approach used here to generate high-performance biomarkers for sex-related BCa progression as well, potentially through the incorporation of multi-omics data and deep learning.

## Methods

### Data Stratification and Cross-Validation Scheme

To elucidate biomarkers associated with gender-specific development and gender-related progression, two analyses were conducted in parallel: one stratified by gender and another stratified by disease status. The gender-stratified study contrasted the expression profiles of tumor tissue versus non-tumor tissue for each gender, while the disease status-stratified study focused on differences between males and females in both diseased and non-tumor tissue. These two stratifications created three core classification tasks: male tumor versus male non-tumor, female tumor versus female non-tumor, and male tumor versus female tumor. The gene feature selection pipeline was set up with 5-fold cross-validation, where the training data for each task was initially divided into 5 equal partitions. Then, the entire pipeline (Figure 7) was executed 5 times, with a different partition acting as the internal validation set in each iteration, while the remaining four folds were used for model training. A cross-validated approach was employed for all three classification tasks to ensure the selected biomarkers were robust. The training data for the sex-specific development and sex-related progression analyses differed slightly. In classifying male tumors versus female tumors, the TCGA-BLCA dataset had a reasonable number of samples for each group: 267 male tumors and 102 female tumors after removing stage I, metastasized, and low-grade samples from consideration (Table 1). However, under 5% of the entire TCGA consisted of non-tumor controls, making the sex-stratified tumor versus non-tumor task impossible to execute. In response, a merged cohort was created to combat this extreme class imbalance. TCGA-BLCA, GTEx, and GSE133624 were independently processed and z-scored before being concatenated together on common genes in the GRCh38.p13 reference genome.

**Figure 7:**
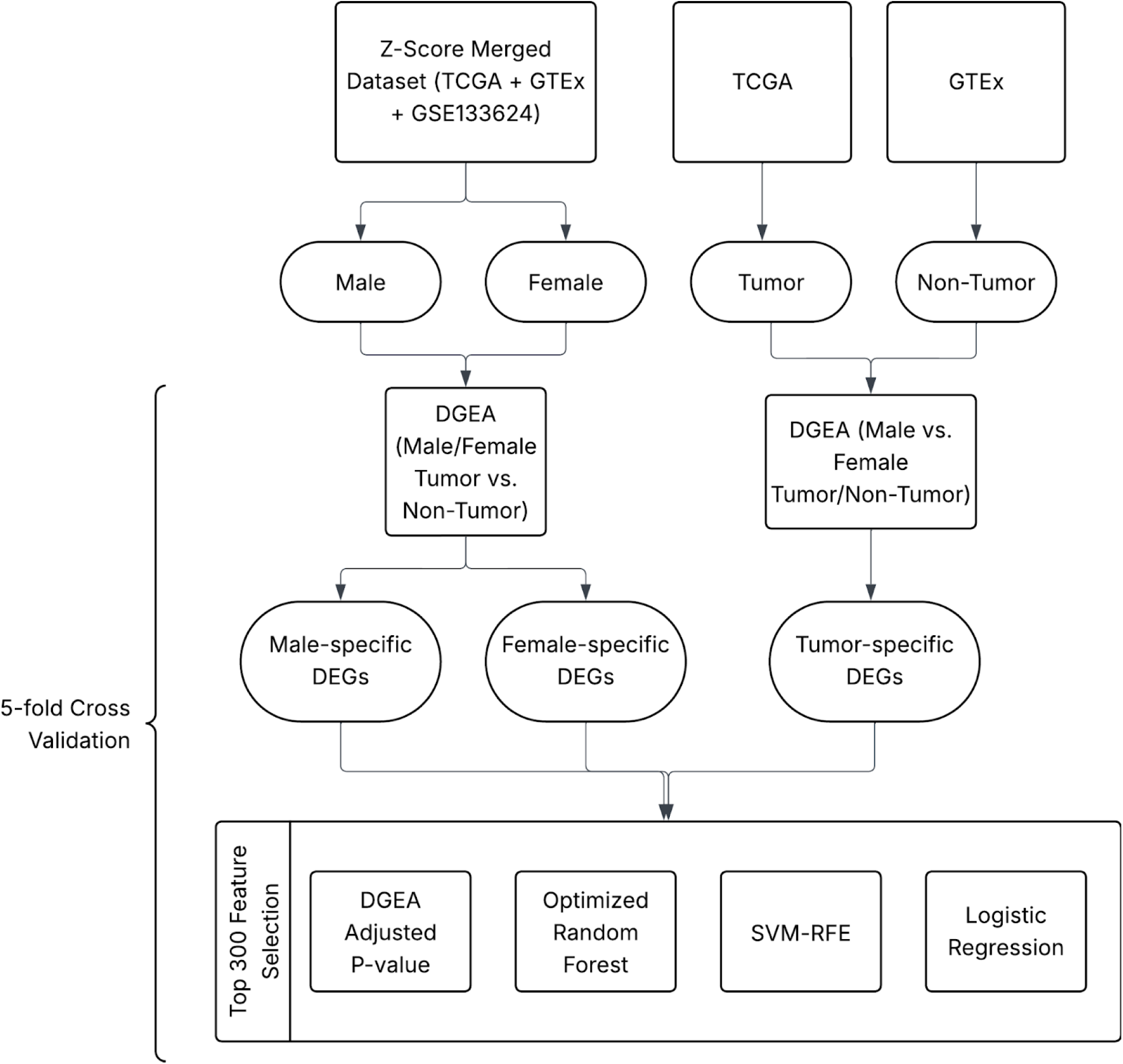
Flowchart of DGEA and feature selection workflow. DEGs specific to differentiating non-tumor males and females are only used to correct for tumor-specific DEGs and do not undergo feature selection.

RNA-seq data from the TCGA-BLCA project were downloaded from the Genomic Data Commons as raw counts. In addition to their gene expressions, clinical information (gender, tumor status, staging, etc.) for each patient was extracted via their patient ID. For the male tumor versus female tumor task, Stage I, low-grade, and metastasized tumors were excluded to ensure gender-related expression differences were not confounded. The male/female tumor vs. non-tumor analyses also incorporated GTEx and NCBI GEO data into this TCGA cohort. Non-tumor bladder tissue raw counts and clinical annotations from the GTEx V10 release were downloaded from the GTEx portal. GSE133624, the final dataset for the merged data pool containing both BCa and paired non-tumor tissue samples, was downloaded from the NCBI GEO as raw counts. For all datasets, Ensembl IDs were used to represent each gene to ensure a common nomenclature before merging. Counts were summed for IDs with multiple entries to avoid duplicate features.

### Dataset Merging and Differential Gene Expression Analysis

To create the merged training dataset, all 3 cohorts were concatenated together on common Ensembl IDs. All genes not appearing across all datasets were excluded from consideration at this point. In the merged data matrix, the original dataset, gender, and disease status for each patient were retained. Afterward, the merged dataset was stratified into male and female subsets, while the TCGA and GTEx datasets acted as the tumor and non-tumor subsets, respectively. Each subset was then split into 5 folds as part of a cross-validation scheme. In this, each fold became the internal validation set while the remaining 4 folds were used for DGEA and feature selection for 5 iterations of the analysis in total for each classification task. The non-tumor subset was an exception to this, as machine learning classification of non-tumor males versus females was not deemed useful. Thus, DGEA for the GTEx data was done outside of the cross-validation scheme to identify DEGs differentiating non-tumor males and females.

All training sets corresponding to each stratified subset were then subjected to DGEA with DESeq2 (59). In the case of the merged training data, the original dataset (batch) was added as a model covariate such that expression differences due to the condition of interest (disease status, gender) were not masked. DEGs between conditions were then selected at a threshold of adjusted p-value < 0.05 and absolute log2 fold change > 1.0. Ensembl IDs were then converted to gene symbols using the GRCh38.p13 reference genome.

### Data Normalization

With DEGs for each stratified subset determined, the merged dataset was first reconstructed. Similarly to how batch was added as a covariate in the DESeq2 model, differences in distributions between datasets had to be accounted for before training feature selection machine learning models. To achieve this, each of the three datasets was independently normalized before concatenation. First, raw counts were converted to TPM values to account for transcript length differences between genes using the GRCh38.p13 reference genome. All TPM values were then subjected to a log2 transformation with a pseudocount of 1 to account for genes with no expression. Afterward, each dataset was independently z-scored to bring the mean to 0 and the standard deviation to 1 for each gene. All 3 normalized datasets were concatenated to create the merged cohort for the male/female tumor versus non-tumor tissue analyses. The efficacy of this correction technique was visualized through UMAP on the data merged with and without prior independent normalization. For the male versus female tumor analysis, the TCGA raw counts were simply converted to TPM values with the GRCh38.p13 reference genome and then subject to a log2 transformation with a pseudocount of 1.

After normalization, the data for all stratified subsets were limited to the previously-identified DEGs. The same 5-fold split applied to the counts data was applied to the normalized TPM training data, so DEGs remained consistent within each fold scheme. However, a key caveat is that DEGs were limited to those unique to each subset. In this, genes differentially expressed between non-tumor and tumor samples for both genders were isolated. Afterward, DEGs with the same direction of regulation for both genders were removed from consideration. For the disease status-stratified analysis, genes differentially expressed between male and female samples for both tumor and non-tumor tissue were excluded from consideration unless oppositely regulated. The purpose of this step was to remove genes that have altered expression in BCa, regardless of gender, and genes with shifted expression between genders that may not be related to BCa. At this point, the data for the non-tumor cohort was deemed obsolete, as its sole purpose was to help remove irrelevant genes from the tumor cohort. As such, only the DEG-limited normalized training data from the male, female, and tumor cohorts moved on to the feature selection phase.

### Feature Selection and Aggregate Panel Construction

Processed training data for each stratified subset was subjected to four different feature selection strategies: DGEA adjusted p-value, optimized RF, SVM-RFE, and cross-validated logistic regression. Mori and colleagues executed 1000 trials of their deep learning-based feature selection on pancreatic cancer gene expression data, tallying how many times each gene appeared in the top 10 and 50 feature importances (60). Olatunji and Cui applied this methodology to an RF model tallying the top 100 genes most important to distant metastasis at each iteration (61). The present approach built upon these implementations by tracking the top 300 genes and summing their RF feature importance values across all 1000 iterations with different random seeds. Before the 1000 iterations, the RF model was hyperparameter optimized with a stratified 5-fold cross-validated exhaustive grid search. The 300 genes with the highest feature importance sums were selected for evaluation for each classification task. For SVM-RFE, a linear kernel was used to utilize variable coefficients as a feature importance metric, and a stratified 5-fold exhaustive search was used to find the optimal value for the regularization parameter. Standardization was applied to each unique training fold within cross-validation rather than the entire inputted subset to avoid data leakage. While slightly stricter than other cancer biomarker studies, 0.1% of uninformative features were removed at each iteration of RFE until 300 remained. For the logistic regression model, a stratified 5-fold exhaustive search identified the optimal regularization parameter value for the L1 penalty, with saga as the solver based on validation performance. As with SVM-RFE, standardization was applied to each unique training fold within cross-validation rather than the entire inputted subset to avoid data leakage. The top 300 genes were selected based on the magnitude of their coefficients. The final feature selection approach extracted the top 300 DEGs by adjusted p-value with an absolute log2 fold change greater than 1.0. For all parameter searches, a composite metric taking the F1 score, the area under the receiver operator curve, and balanced accuracy into account was utilized to adjust for class imbalance (0.5*F1 + 0.3*AUROC + 0.2* balanced accuracy). The F1 score was weighted the most heavily due to its sensitivity to the true positive rate and established utility in imbalanced human health datasets (62). AUROC is particularly relevant to diagnostic problems and is also robust to class imbalance, but is less established compared to the F1 score (22,63). Finally, balanced accuracy was included with the smallest weight to represent a simpler measure of model performance. All machine learning models, feature selection approaches, and hyperparameter optimization searches were implemented with the scikit-learn package. The result of feature selection was four top 300-gene panels for each of the three cohorts within each cross-validation scheme.

Across 5 fold schemes and four feature selection methods, 20 top 300-gene panels were generated for each classification task. All rankings within a task were combined into an aggregated top 300 panel using RRA to synthesize findings across all feature selection approaches and combinations of training folds (64). This assigned a score to each gene, with lower scores being assigned to those highly ranked across all lists. To visualize the RRA scores, a negative log10 transformation was applied to the scores of the top genes and a bar graph was constructed with the matplotlib package. Consensus top 300 panels for each selection method were also produced, aggregating the five rankings produced by each fold scheme. The overlap in genes between the consensus panels for each feature selection approach was demonstrated by creating a Venn diagram for each classification task using the pyvenn package. The four cross-fold consensus panels and the aggregated ranking were then considered for validation on external datasets for each classification task (Figure 8).

**Figure 8:**
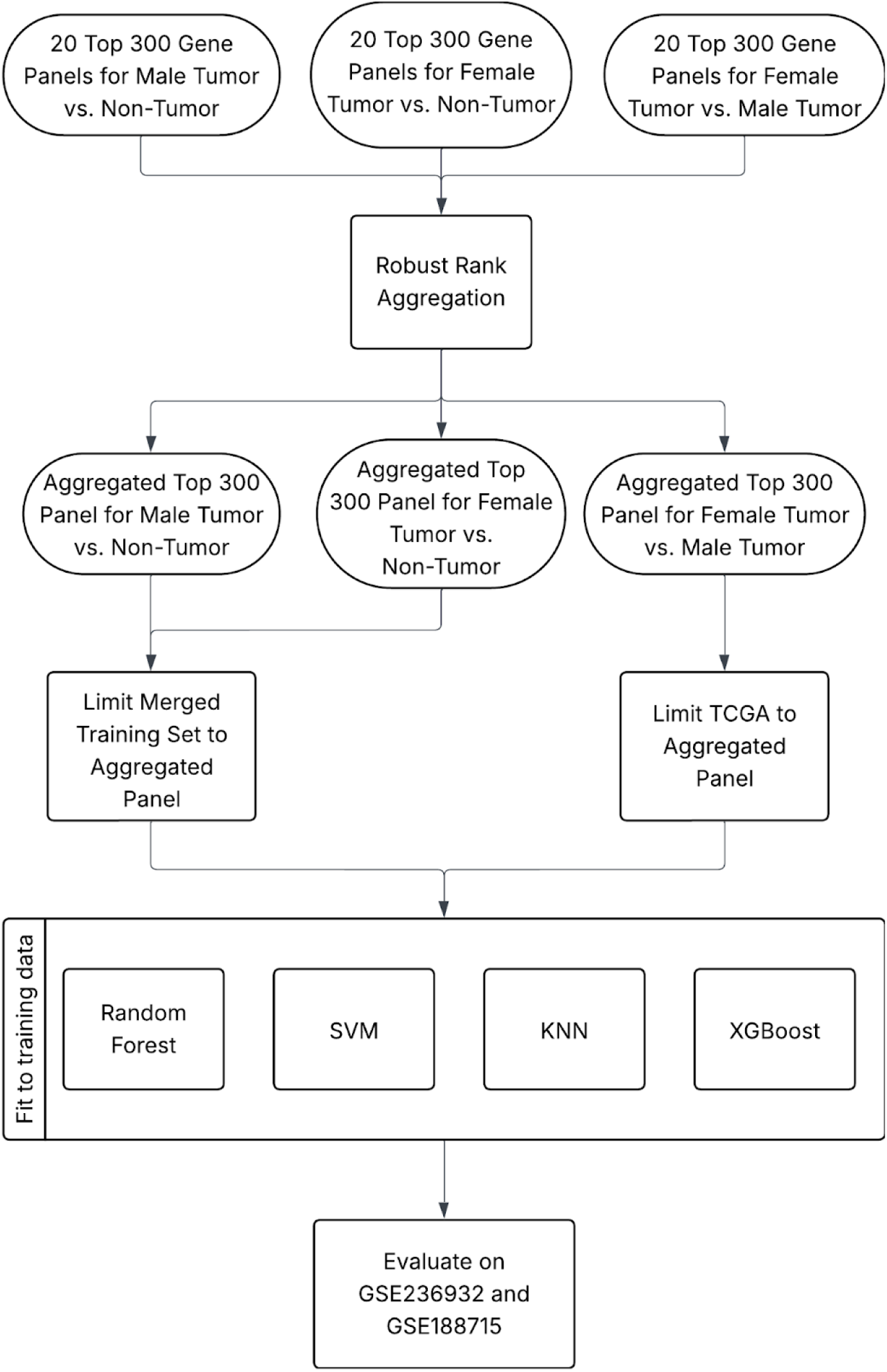
Flowchart of evaluation workflow. In the case of the merged training set for male/female tumor versus non-tumor, GSE236932 and GSE188715 were z-scored before scoring.

### Gene Panel Evaluation

To evaluate each gene panel, four evaluation models were trained: RF, SVM with linear kernel, KNN classifier, and XGBoost. XGBoost was outside of scikit-learn and required the installation of the xgboost module. For each, the classifier was introduced to the training data and then applied to the external testing sets, all limited to the gene panel of interest. The external cohorts consisted of RNA-seq NCBI GEO datasets GSE236932 and GSE188715, whose expressions were downloaded as TPM values calculated with GRCh38.p13. Genes with more than one entry had their expressions averaged, and the same log2 transformation with a pseudocount of 1 was applied, just as was done for the training data. For the male/female tumor versus non-tumor tasks, each external dataset was z-scored independently of the training data to bring them to the same scale. Ideally, the mean and standard deviation from the training set would be applied to the test data, but this was not possible due to standardization being done independently for each of the three training datasets prior to merging.

Each top 300 gene panel was further subsetted (top 300, 250, 200, etc.) to judge whether diminishing the feature count enhanced predictive ability. Before evaluating the classifiers on the test set, their hyperparameters were optimized on the training data using 5-fold cross-validation with the previously mentioned composite metric. For the distance-based models (SVM and KNN), the classifiers were pipelined with a standardization step to prevent data leakage during cross-validation. Due to their smaller hyperparameter spaces, exhaustive search was appropriate for SVM and KNN classifiers. For RF and XGBoost, a randomized grid search with 200 iterations and a Bayesian search implementation from scikit-optimize with 50 iterations were required to explore the much larger number of possible hyperparameter combinations. The performance of each subset on the internal and external testing sets was compared using accuracy, F1 score, AUROC, and the weighted composite metric. ROC curves for each feature subset in each external dataset were produced using the RF, SVM, and XGBoost evaluation models, and those with the top AUROC scores were visualized together. Such a detailed evaluation of each top 300 gene panel identified the selection method and subset with the best classification performance for each task.

### Pathway Enrichment and PPI Network Analysis

Pathway enrichment was carried out on the aggregated top 300 panel for each classification task using clusterProfiler from Bioconductor (65). Significant ontologies were identified as those having a q-value less than 0.05 and the KEGG pathways and GO biological process databases were utilized. However, at this threshold the male and female tumor versus non-tumor panels did not yield any significant hits, so the query was expanded to the top 500 genes for each task. Dot plots were constructed with the ggplot2 graphical interface to represent the top 10 pathways for each task/database combination.

PPI network analysis was performed by inputting the top 300 genes of each aggregated panel into the STRING database web interface to generate an edge-node graph of protein interactions. A TSV of node relationships was outputted and all gene products without edges or a combined score less than 0.4 were excluded from further analysis. All three networks were reconstructed by importing the STRING TSV into Cytoscape. Within Cytoscape, MCODE was used to identify significant gene modules while cytoHubba helped to execute topological analysis and isolate important nodes. For MCODE, a degree cutoff of 2, node score cutoff of 0.2, k-core of 2, and max depth of 100 were utilized with the haircut setting turned on. Significant clusters were defined as those having a score of 3 or greater. For cytoHubba, the top 10 genes in terms of maximal clique centrality were determined for each of the three networks and classified as hub genes. Pathway enrichment was performed for gene modules containing hub genes using the Enrichr web interface to obtain significant terms in the Reactome database (66). All pathways with a q-value of 0.05 or less were considered.

## Acknowledgements

We are grateful to the Thomas H. Gosnell School of Life Sciences and the College of Science at RIT for administrative and financial support.

## Availability of methods

The code used to generate the results in this study can be found on GitHub: https://github.com/rit-cui-lab/Machine-learning-based-determination-of-sex-related-bladder-cancer-biomarkers

## References

1. SEER [Internet]. [cited 2025 Dec 19]. Cancer of the Urinary Bladder - Cancer Stat Facts. Available from: https://seer.cancer.gov/statfacts/html/urinb.html

2. Jakus D, Šolić I, Jurić I, Borovac JA, Šitum M. The Impact of the Initial Clinical Presentation of Bladder Cancer on Histopathological and Morphological Tumor Characteristics. J Clin Med. 2023 June 25;12(13):4259.

3. Lopez-Beltran A, Cookson MS, Guercio BJ, Cheng L. Advances in diagnosis and treatment of bladder cancer. BMJ. 2024 Feb 12;384:e076743.

4. Flores Monar GV, Reynolds T, Gordon M, Moon D, Moon C. Molecular Markers for Bladder Cancer Screening: An Insight into Bladder Cancer and FDA-Approved Biomarkers. Int J Mol Sci. 2023 Sept 21;24(18):14374.

5. Siegel RL, Kratzer TB, Giaquinto AN, Sung H, Jemal A. Cancer statistics, 2025. Ca. 2025;75(1):10–45.

6. Dai X, Gakidou E, Lopez AD. Evolution of the global smoking epidemic over the past half century: strengthening the evidence base for policy action. Tob Control. 2022 Mar 1;31(2):129–37.

7. Poli C, Trétarre B, Trouche-Sabatier S, Foucan AS, Abdo N, Poinas G, et al. Sex differences in muscle-invasive bladder cancers: A study of a French regional population. Fr J Urol. 2025 Jan 1;35(1):102723.

8. Ark JT, Alvarez JR, Koyama T, Bassett JC, Blot WJ, Mumma MT, et al. Variation in the diagnostic evaluation among persons with hematuria: influence of gender, race, and risk factors for bladder cancer. J Urol. 2017 Nov;198(5):1033–8.

9. Radkiewicz C, Edgren G, Johansson ALV, Jahnson S, Häggström C, Akre O, et al. Sex Differences in Urothelial Bladder Cancer Survival. Clin Genitourin Cancer. 2020 Feb 1;18(1):26–34.e6.

10. Li P, Chen J, Miyamoto H. Androgen Receptor Signaling in Bladder Cancer. Cancers. 2017 Feb 22;9(2):20.

11. Kolyvas EA, Caldas C, Kelly K, Ahmad SS. Androgen receptor function and targeted therapeutics across breast cancer subtypes. Breast Cancer Res. 2022 Nov 14;24(1):79.

12. Goto T, Yasui M, Teramoto Y, Nagata Y, Mizushima T, Miyamoto H. Latrophilin-3 as a downstream effector of the androgen receptor induces urothelial tumorigenesis. Mol Carcinog. 2024;63(10):1847–54.

13. Inoue S, Ide H, Mizushima T, Jiang G, Kawahara T, Miyamoto H. ELK1 promotes urothelial tumorigenesis in the presence of an activated androgen receptor. Am J Cancer Res. 2018 Nov 1;8(11):2325–36.

14. Elahi Najafi Ma, Matsukawa T, Miyamoto H. Recent advances in understanding the role of sex hormone receptors in urothelial cancer. Oncol Res. 33(6):1255–70.

15. Doshi B, Athans SR, Woloszynska A. Biological differences underlying sex and gender disparities in bladder cancer: current synopsis and future directions. Oncogenesis. 2023 Sept 4;12(1):44.

16. Fuentes N, Silveyra P. Estrogen receptor signaling mechanisms. Adv Protein Chem Struct Biol. 2019;116:135–70.

17. Goto T, Miyamoto H. The Role of Estrogen Receptors in Urothelial Cancer. Front Endocrinol. 2021 Mar 16;12:643870.

18. Tatenuma T, Matsukawa T, Goto T, Jiang G, Sharma A, Najafi MAE, et al. GULP1 as a Downstream Effector of the Estrogen Receptor-β Modulates Cisplatin Sensitivity in Bladder Cancer. Cancer Genomics Proteomics. 2024 Nov 1;21(6):557–65.

19. Koti M, Ingersoll MA, Gupta S, Lam CM, Li X, Kamat AM, et al. Sex Differences in Bladder Cancer Immunobiology and Outcomes: A Collaborative Review with Implications for Treatment. Eur Urol Oncol. 2020 Oct 1;3(5):622–30.

20. Qiu H, Makarov V, Bolzenius JK, Halstead A, Parker Y, Wang A, et al. KDM6A Loss Triggers an Epigenetic Switch That Disrupts Urothelial Differentiation and Drives Cell Proliferation in Bladder Cancer. Cancer Res. 2023 Mar 15;83(6):814–29.

21. Wang Y, Bhandary P, Griffin K, Moore JH, Li X, Wang ZP. Integrative multi-omics study identifies sex-specific molecular signatures and immune modulation in bladder cancer. Front Bioinforma [Internet]. 2025 May 19 [cited 2025 Nov 16];5. Available from: https://www.frontiersin.org/journals/bioinformatics/articles/10.3389/fbinf.2025.1575790/full

22. Çorbacıoğlu ŞK, Aksel G. Receiver operating characteristic curve analysis in diagnostic accuracy studies: A guide to interpreting the area under the curve value. Turk J Emerg Med. 2023 Oct 3;23(4):195–8.

23. Low JY, Esopi DM, Lim Y, Vaghasia AM, Tsai H, Zheng Q, et al. Abstract B011: PRAC1 epigenetic silencing in castration resistant prostate cancer and its novel role in androgen receptor biology. Cancer Res. 2023 June 2;83(11_Supplement):B011.

24. Kim YW, Yoon HY, Seo SP, Lee SK, Kang HW, Kim WT, et al. Clinical Implications and Prognostic Values of Prostate Cancer Susceptibility Candidate Methylation in Primary Nonmuscle Invasive Bladder Cancer. Dis Markers. 2015;2015:402963.

25. Mo XC, Zhang ZT, Song MJ, Zhou ZQ, Zeng JX, Du YF, et al. Screening and identification of hub genes in bladder cancer by bioinformatics analysis and KIF11 is a potential prognostic biomarker. Oncol Lett. 2021 Mar;21(3):205.

26. Xu Y, Wu G, Li J, Li J, Ruan N, Ma L, et al. Screening and Identification of Key Biomarkers for Bladder Cancer: A Study Based on TCGA and GEO Data. BioMed Res Int. 2020 Jan 23;2020:8283401.

27. Terry S, Queires L, Gil-Diez-de-Medina S, Chen MW, Taille A de la, Allory Y, et al. Protocadherin-PC Promotes Androgen-Independent Prostate Cancer Cell Growth. The Prostate. 2006 July 1;66(10):1100–13.

28. Chen F, Li Z, Deng C, Yan H. Integrated analysis identifying new lncRNA markers revealed in ceRNA network for tumor recurrence in papillary thyroid carcinoma and build of nomogram. J Cell Biochem. 2019;120(12):19673–83.

29. Zhang S, Chen R. LINC01140 regulates osteosarcoma proliferation and invasion by targeting the miR-139-5p/HOXA9 axis. Biochem Biophys Rep. 2022 June 28;31:101301.

30. Zhao L, Wang G, Qi H, Yu L, Yin H, Sun R, et al. LINC00330/CCL2 axis-mediated ESCC TAM reprogramming affects tumor progression. Cell Mol Biol Lett. 2024 May 20;29:77.

31. Zhu H, Yu J, Zhu H, Guo Y, Feng S. Identification of key lncRNAs in colorectal cancer progression based on associated protein–protein interaction analysis. World J Surg Oncol. 2017 Aug 10;15:153.

32. Frenay J, Bellaye PS, Oudot A, Helbling A, Petitot C, Ferrand C, et al. IL-1RAP, a Key Therapeutic Target in Cancer. Int J Mol Sci. 2022 Nov 29;23(23):14918.

33. Han J, Wang J zi, Yang X, Yu H, Zhou R, Lu HC, et al. METTL3 promote tumor proliferation of bladder cancer by accelerating pri-miR221/222 maturation in m6A-dependent manner. Mol Cancer. 2019 June 22;18:110.

34. Cucu D. The Potential of TRPA1 as a Therapeutic Target in Cancer—A Study Using Bioinformatic Tools. Pharmaceuticals. 2024 Dec 9;17(12):1657.

35. He RQ, Huang ZG, Zhai GQ, Huang SN, Gu YY, Chen G, et al. Small Nucleolar RNAs (snoRNAs)-Based Risk Score Classifier Predicts Overall Survival in Bladder Carcinoma. Med Sci Monit Int Med J Exp Clin Res. 2020 Oct 26;26:e926273-1-e926273-14.

36. Jin Z, Yao J, Xie N, Cai L, Qi S, Zhang Z, et al. Melittin Constrains the Expression of Identified Key Genes Associated with Bladder Cancer. J Immunol Res. 2018 May 3;2018:5038172.

37. Wang W, Gao Y, Liu Y, Xia S, Xu J, Qin L, et al. Pan-cancer analysis reveals MTTP as a prognostic and immunotherapeutic biomarker in human tumors. Front Immunol. 2025 Mar 27;16:1549965.

38. Piazza GA, Ward A, Chen X, Maxuitenko Y, Coley A, Aboelella NS, et al. PDE5 and PDE10 inhibition activates cGMP/PKG signaling to block Wnt/β-catenin transcription, cancer cell growth, and tumor immunity. Drug Discov Today. 2020 Aug 1;25(8):1521–7.

39. Lan T, Li Y, Wang Y, Wang ZC, Mu CY, Tao AB, et al. Increased endogenous PKG I activity attenuates EGF-induced proliferation and migration of epithelial ovarian cancer via the MAPK/ERK pathway. Cell Death Dis. 2023 Jan 19;14(1):39.

40. Liu Z, Zhang X, Ben T, Li M, Jin Y, Wang T, et al. Focal adhesion in the tumour metastasis: from molecular mechanisms to therapeutic targets. Biomark Res. 2025 Mar 5;13(1):38.

41. Knights AJ, Funnell APW, Crossley M, Pearson RCM. Holding Tight: Cell Junctions and Cancer Spread. Trends Cancer Res. 2012;8:61–9.

42. Leylek TR, Jeusset LM, Lichtensztejn Z, McManus KJ. Reduced Expression of Genes Regulating Cohesion Induces Chromosome Instability that May Promote Cancer and Impact Patient Outcomes. Sci Rep. 2020 Jan 17;10(1):592.

43. Zhang W, Wang Y, Tang Q, Li Z, Sun J, Zhao Z, et al. PAX2 mediated upregulation of ESPL1 contributes to cisplatin resistance in bladder cancer through activating the JAK2/STAT3 pathway. Naunyn Schmiedebergs Arch Pharmacol. 2024 Sept 1;397(9):6889–901.

44. Chi M, Liu J, Mei C, Shi Y, Liu N, Jiang X, et al. TEAD4 functions as a prognostic biomarker and triggers EMT via PI3K/AKT pathway in bladder cancer. J Exp Clin Cancer Res. 2022 May 17;41(1):175.

45. Miyamoto H, Yao JL, Chaux A, Zheng Y, Hsu I, Izumi K, et al. Expression of androgen and oestrogen receptors and its prognostic significance in urothelial neoplasm of the urinary bladder. BJU Int. 2012;109(11):1716–26.

46. Sikic D, Wirtz RM, Wach S, Dyrskjøt L, Erben P, Bolenz C, et al. Androgen Receptor mRNA Expression in Urothelial Carcinoma of the Bladder: A Retrospective Analysis of Two Independent Cohorts. Transl Oncol. 2019 Mar 2;12(4):661–8.

47. Hu J, Zhang J, Han B, Qu Y, Zhang Q, Yu Z, et al. PLXNA1 confers enzalutamide resistance in prostate cancer via AKT signaling pathway. Neoplasia N Y N. 2024 Sept 2;57:101047.

48. Zhou C, Zhang X, Ma H, Zhou Y, Meng Y, Chen C, et al. USP54 is a potential therapeutic target in castration-resistant prostate cancer. BMC Urol. 2024 Feb 6;24(1):32.

49. Qiu D, Hu J, Hu J, Yu A, Othmane B, He T, et al. PMEPA1 Is a Prognostic Biomarker That Correlates With Cell Malignancy and the Tumor Microenvironment in Bladder Cancer. Front Immunol. 2021 Oct 28;12:705086.

50. He Z, Wang C, Xue H, Zhao R, Li G. Identification of a Metabolism-Related Risk Signature Associated With Clinical Prognosis in Glioblastoma Using Integrated Bioinformatic Analysis. Front Oncol. 2020 Sept 3;10:1631.

51. Huang J, Zhang L, Wan D, Zhou L, Zheng S, Lin S, et al. Extracellular matrix and its therapeutic potential for cancer treatment. Signal Transduct Target Ther. 2021 Apr 23;6:153.

52. Necula L, Matei L, Dragu D, Pitica I, Neagu A, Bleotu C, et al. Collagen Family as Promising Biomarkers and Therapeutic Targets in Cancer. Int J Mol Sci. 2022 Jan;23(20):12415.

53. Zeng XT, Liu XP, Liu TZ, Wang XH. The clinical significance of COL5A2 in patients with bladder cancer. Medicine (Baltimore). 2018 Mar 9;97(10):e0091.

54. Darb-Esfahani S, Kronenwett R, von Minckwitz G, Denkert C, Gehrmann M, Rody A, et al. Thymosin beta 15A (TMSB15A) is a predictor of chemotherapy response in triple-negative breast cancer. Br J Cancer. 2012 Nov;107(11):1892–900.

55. Shabbir R, Quiles CG, Lane B, Zeef L, Hoskin PJ, Choudhury A, et al. Gene Expression in Muscle-Invasive and Non-Muscle-Invasive Bladder Cancer Cells Exposed to Hypoxia. Cancers. 2025 Aug 11;17(16):2624.

56. Shen L, Zhang J, Zheng Z, Yang F, Liu S, Wu Y, et al. PHGDH Inhibits Ferroptosis and Promotes Malignant Progression by Upregulating SLC7A11 in Bladder Cancer. Int J Biol Sci. 2022 Aug 29;18(14):5459–74.

57. da Silva IM, Maraslis FT, Kawasaki JAI, Aida NK, Barcelos GRM, Koike A, et al. Allelic variants in xenobiotic metabolism genes predict susceptibility and worse prognosis of urothelial bladder cancer. Pathol - Res Pract. 2025 Feb 1;266:155767.

58. Garg M, Maurya N. WNT/β-catenin signaling in urothelial carcinoma of bladder. World J Nephrol. 2019 Sept 26;8(5):83–94.

59. Love MI, Huber W, Anders S. Moderated estimation of fold change and dispersion for RNA-seq data with DESeq2. Genome Biol. 2014 Dec 5;15(12):550.

60. Mori Y, Yokota H, Hoshino I, Iwatate Y, Wakamatsu K, Uno T, et al. Deep learning-based gene selection in comprehensive gene analysis in pancreatic cancer. Sci Rep. 2021 Aug 13;11:16521.

61. Olatunji I, Cui F. Multimodal AI for prediction of distant metastasis in carcinoma patients. Front Bioinforma. 2023 May 9;3:1131021.

62. Diallo R, Edalo C, Awe OO. Machine Learning Evaluation of Imbalanced Health Data: A Comparative Analysis of Balanced Accuracy, MCC, and F1 Score. In: Awe OO, A. Vance E, editors. Practical Statistical Learning and Data Science Methods: Case Studies from LISA 2020 Global Network, USA [Internet]. Cham: Springer Nature Switzerland; 2025 [cited 2025 Dec 19]. p. 283–312. Available from: 10.1007/978-3-031-72215-8_12

63. Richardson E, Trevizani R, Greenbaum JA, Carter H, Nielsen M, Peters B. The receiver operating characteristic curve accurately assesses imbalanced datasets. Patterns. 2024 June 14;5(6):100994.

64. Kolde R, Laur S, Adler P, Vilo J. Robust rank aggregation for gene list integration and meta-analysis. Bioinformatics. 2012 Feb 15;28(4):573–80.

65. Yu G, Wang LG, Han Y, He QY. clusterProfiler: an R Package for Comparing Biological Themes Among Gene Clusters. OMICS J Integr Biol. 2012 May;16(5):284–7.

66. Xie Z, Bailey A, Kuleshov MV, Clarke DJB, Evangelista JE, Jenkins SL, et al. Gene Set Knowledge Discovery with Enrichr. Curr Protoc. 2021;1(3):e90.

